# Instability in 18S and 5.8S rDNA copy numbers and 18S:5.8S ratios in longitudinally-collected human DNA samples detected by monochrome multiplex qPCR

**DOI:** 10.1101/361840

**Authors:** Chris B. Ligerman, Lisa Baird, Mark F. Leppert, Richard M. Cawthon

**Affiliations:** Dept. of Human Genetics, University of Utah, Salt Lake City, Utah, United States of America

## Abstract

Ribosomal DNA encodes the structural RNAs of the ribosomes. Ribosomal DNA instability is a major contributor to aging in yeast, but its role in human aging and longevity is largely unknown. Human 45S rDNA encodes the 18S, 5.8S, and 28S ribosomal RNAs; ranges in copy number from 60 to > 800 per cell, distributed as tandem repeats along the short arms of the five acrocentric chromosomes (p13, p14, p15, p21, and p22); and is prone to frequent homologous and non-homologous recombination. Here we present two multiplex quantitative PCR assays, one for 18S rDNA normalized to the single copy gene beta-globin (HBB), and the other for 5.8S rDNA normalized to the single copy gene albumin (ALB). Longitudinally-collected pairs of DNAs from bloods drawn approximately 16 years apart from 40 females and 39 males, aged < 1 to 77 years at first blood draw, from the Utah CEPH families were assayed. Ribosomal DNA copy number varied over a four-fold range between subjects and increased approximately 14% across the lifespan. Longitudinal within-individual gains in copy number up to +68% and losses down to ‐25% were observed, while repeated assays of single DNA samples varied approximately +/- 10%. While 18S and 5.8S rDNA tended to be gained and lost together, the 18S:5.8S ratio was also unstable longitudinally, with increases up to +19% and decreases down to ‐15% observed. The 18S:5.8S ratio at second draw, relative to that at first draw, increased significantly across the lifespan in males, but not in females. To our knowledge this is the first report of within-individual longitudinal changes in the human 18S and 5.8S rDNA copy numbers. These assays will facilitate investigations of the biology of ribosomal RNA genes and their roles in health and disease across the human lifespan.

## Introduction

The role of rDNA instability in human aging and longevity is largely unknown. The copy number of rDNA (the genes encoding the ribosomal RNAs) is unstable in eukaryotes [1]. Stabilizing the rDNA copy number in yeast extends lifespan [2]. Human 45S rDNA is tandemly repeated along the p-arms of the acrocentric chromosomes 13, 14, 15, 21, and 22. Each 45S rDNA repeat typically contains one copy each of the 18S, 5.8S, and 28S rDNA genes. The average 45S rDNA copy number per cell varies more than 10-fold between individuals [3], can change significantly between parents and offspring [4], and is altered in various disease conditions (e.g. cancer [5] and neurodegeneration [6]). Evidence that the components of the 18S-5.8S-28S rDNA repeating unit can become unstable and deviate from their expected 1:1:1 ratio was provided by a quantitative PCR study that found lower copy numbers of 5.8S and 28S rDNA relative to 18S rDNA, in the adipose tissue of older individuals vs. young individuals [7]; and a full genome sequencing analysis of human lymphoblastoid cell line DNAs that found fewer sequence reads for 5.8S and 28S rDNA than for 18S rDNA, with this deviation being more pronounced in men than in women [8]. Here we present two novel monochrome multiplex qPCR assays to make copy number assessments of the 18S and 5.8S rDNA amenable to low cost, high-throughput screening, and we report the first longitudinal study of human rDNA copy number instability, in pairs of DNA samples extracted from blood draws collected several years apart from the same individuals.

## Results

### 18S and 5.8S rDNA copy numbers are highly correlated in whole blood DNA samples, and vary between individuals over an approximately 4-fold range

The 79 subjects in this study were selected from the 3-generation Utah CEPH families (generation I, ages 57-77 years at first blood draw in the early 1980s; generation II, ages 35-59; generation III, ages <1-28) so that within each generation, all individuals of the same sex were unrelated (from different families). The interval between within-individual blood draws ranged from 12.5 to 20.3 years. The primers and conditions for the monochrome multiplex qPCR reactions are given in the Methods section. The 18S rDNA primer pair and the HBB (beta-globin) primer pair were used for one multiplex assay, and the 5.8S rDNA primer pair and ALB (albumin) primer pair were used for the second multiplex assay. HBB and ALB are single copy genes; normalizing to their amplification signals serves as a proxy for normalizing to cell count. We used the relative qPCR method, where the results for each DNA sample in the study are expressed relative to those for a reference DNA sample, using the Standard Curve method.

Fig 1 plots the ratio of the 18S rDNA copy number to the HBB copy number (x axis) vs. the ratio of the 5.8S rDNA copy number to the ALB copy number (y axis). These ratios are relative to those in the reference DNA, in which each ratio is assigned the value of 1.00. These results are consistent with the 18S to 5.8S rDNA copy number ratio being *approximately* 1:1 in all the DNA samples tested, although subsequent analyses revealed significant deviations from 1:1 (see below). In absolute numbers, based on the differences in crossing thresholds for the rDNAs vs. those for the single copy genes, we estimate that the average rDNA copy number per cell in these subjects ranges from ~170 – 740; however, absolute quantification of copy numbers by qPCR will require the use of reference samples with known copy numbers of the targeted templates, which was not done for this relative quantification study.

**Fig 1.**
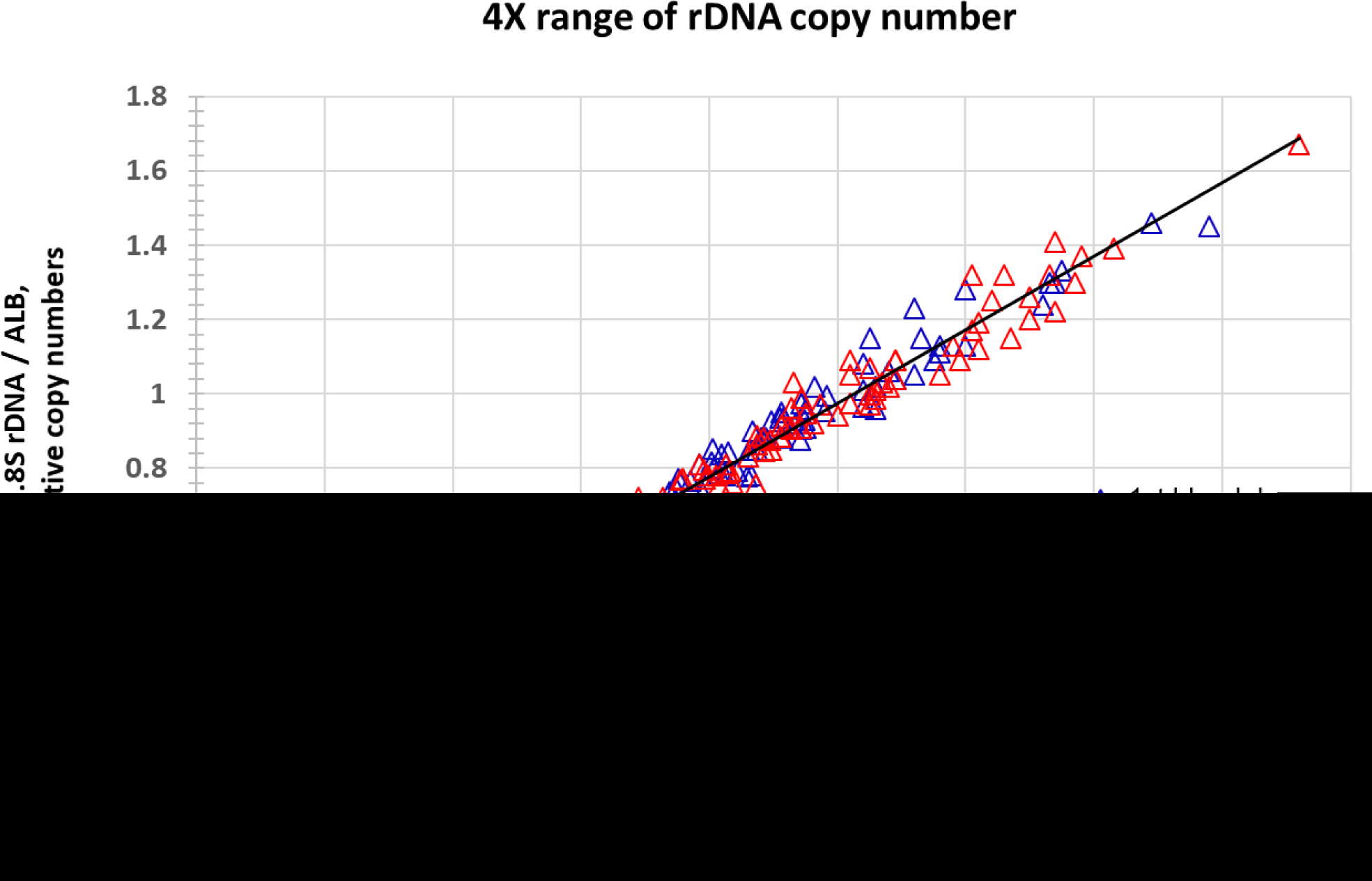
Relative quantitative PCR determination of the average rDNA copy number per cell in 79 individuals, as compared to that in a reference DNA sample. The 18S rDNA is normalized to the single copy gene HBB (x axis), and the 5.8S rDNA is normalized to the single copy gene ALB (y axis). Blue triangles, 1st blood draw DNA samples. Red triangles, 2nd blood draw DNAs.

### Change in 18S and 5.8S rDNA copy numbers cross-sectionally, and longitudinally

Fig 2, top panel plots age at blood draw vs. the copy numbers of 18S and 5.8S rDNA (relative to those in the reference DNA) for all females in the study for each of the two blood draws. There is a trend toward increasing 18S and 5.8S rDNA copy numbers with increasing age, for both the first and second blood draws. First draw rDNA copy numbers increased approximately 1.29% per decade (blue regression line), and second draw rDNA copy numbers increased approximately 1.47% per decade (red regression line). The middle panel shows that both gains and losses of rDNA copies occur across the lifespan, with gains in copy number more common than losses, particularly after age 60 years. The bottom panel shows that 18S copies and 5.8S copies tend to be gained and lost together. Within-individual longitudinal gains in rDNA copy number up to +68% and losses down to ‐25% were observed in these subjects, while repeated assays of single DNA samples varied approximately +/- 10%.

**Fig 2.**
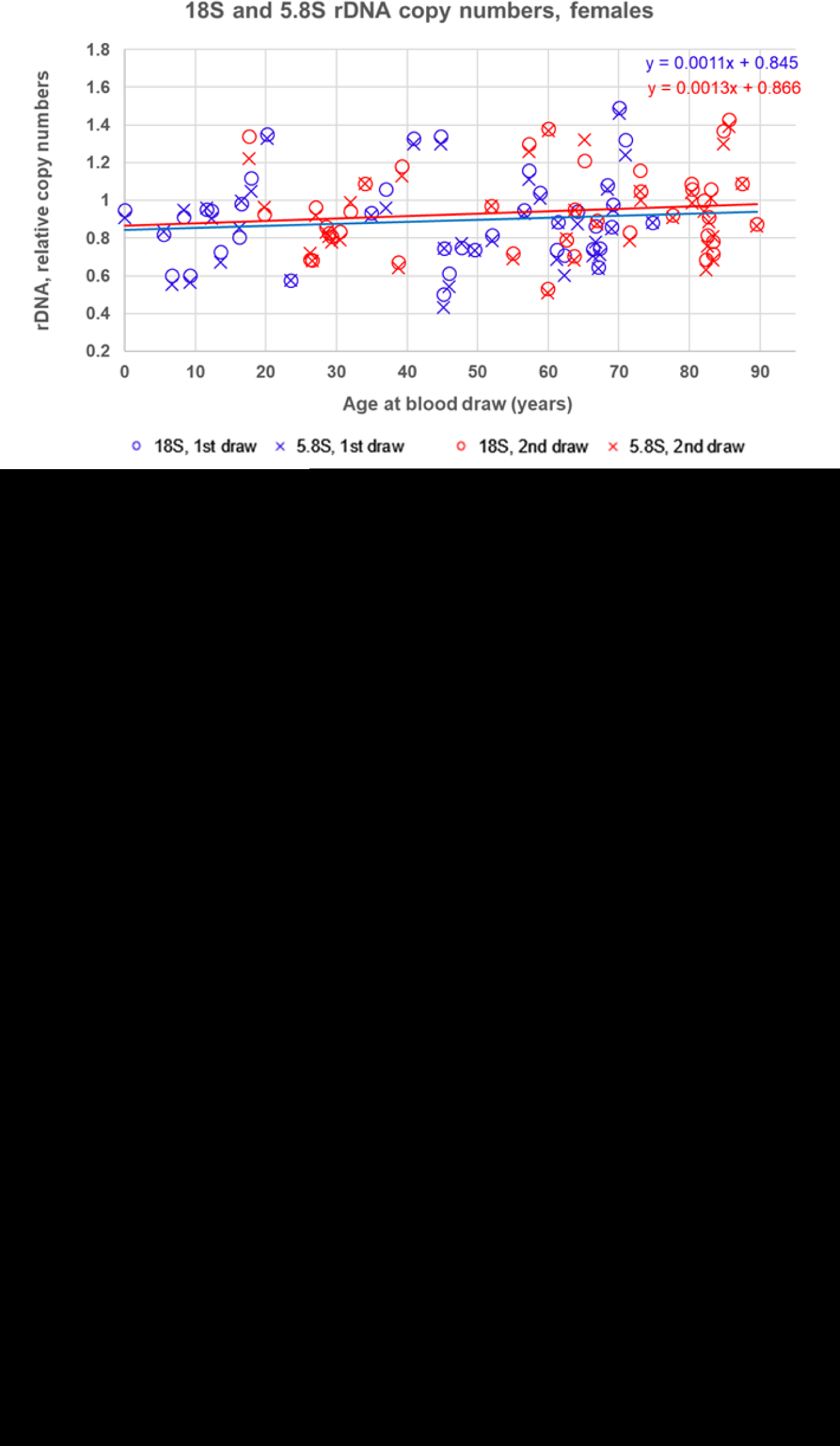
Average 18S and 5.8S copy numbers per cell in 40 female subjects, relative to those in a reference DNA sample, in pairs of blood samples drawn years apart, and longitudinal change in copy number between blood draws. Top panel, age at blood draw vs. relative rDNA/single copy gene ratio. The solid blue line shows the linear regression of age vs. the average of the 18S and 5.8S ratio for each subject, at first blood draw. The solid red line shows the linear regression of age vs. the average of the 18S and 5.8S ratio for each subject, at second blood draw. Middle panel, age at first blood draw vs. the percentage change in the rDNA copy number between blood draws. The interval between blood draws ranged from 12.5 to 20.1 years. Bottom panel, the percentage change in the 18S rDNA copy number between blood draws vs. the percentage change in the 5.8S rDNA copy number in the same DNA samples.

Fig 3 shows analyses for all males in the study, equivalent to those shown in Fig 2 for females, with results very similar to those found in females. The top panel shows that in the males first draw rDNA copy numbers increased approximately 2.05% per decade (blue regression line), and second draw rDNA copy numbers increased approximately 3.14% per decade (red regression line).

**Fig 3.**
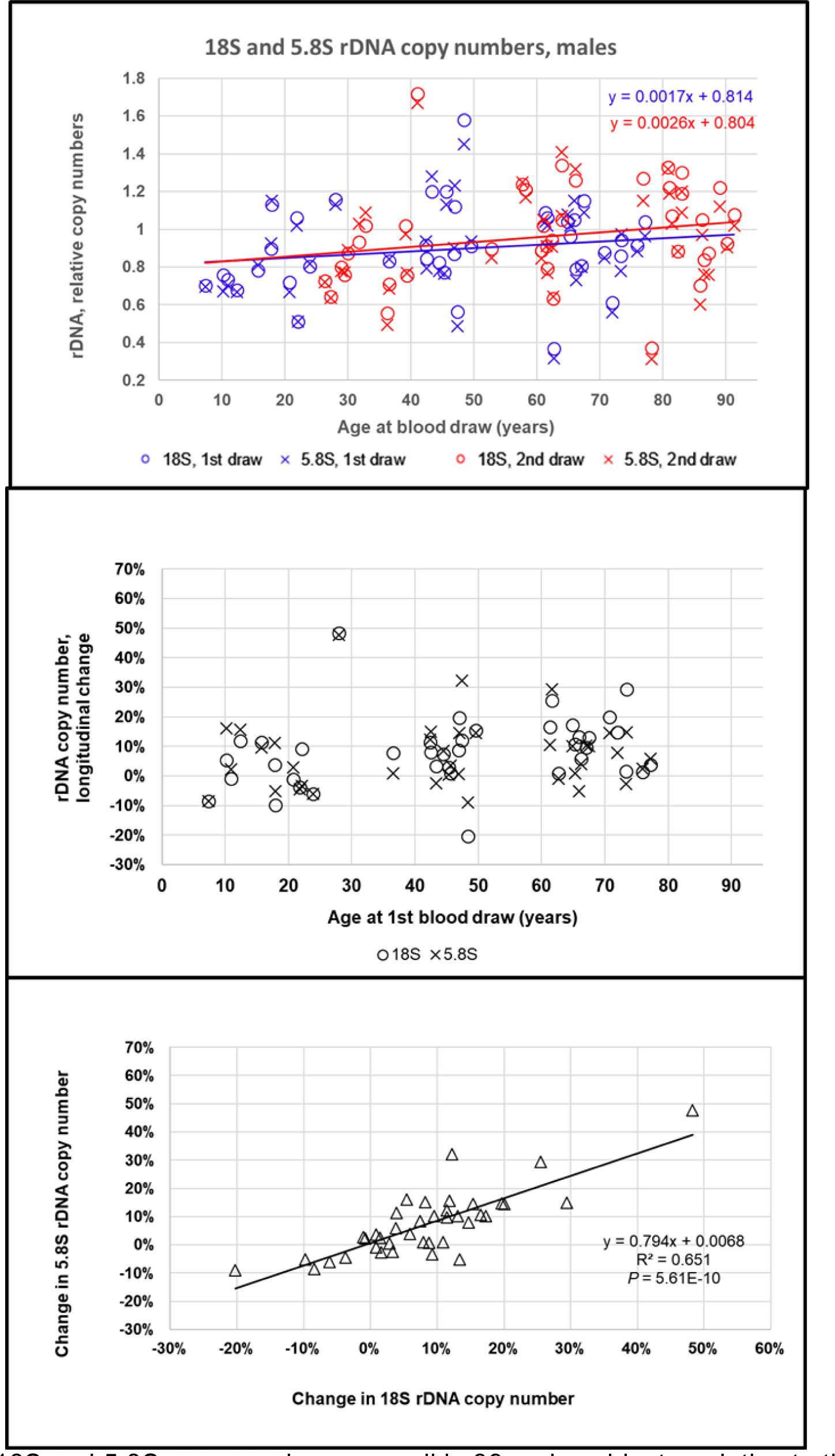
Average 18S and 5.8S copy numbers per cell in 39 male subjects, relative to those in a reference DNA sample, in pairs of blood samples drawn years apart, and longitudinal change in copy number between blood draws. Top panel, age at blood draw vs. relative rDNA/single copy gene ratio. The solid blue line shows the linear regression of age vs. the average of the 18S and 5.8S ratio for each subject, at first blood draw. The solid red line shows the linear regression of age vs. the average of the 18S and 5.8S ratio for each subject, at second blood draw. Middle panel, age at first blood draw vs. the percentage change in the rDNA copy number between blood draws. The interval between blood draws ranged from 12.5 to 20.3 years. Bottom panel, the percentage change in the 18S rDNA copy number between blood draws vs. the percentage change in the 5.8S rDNA copy number in the same DNA samples.

### Instability of the 18S:5.8S rDNA copy number ratio in cross-sectional data

Fig 4, top panel shows a trend toward an increasing 18S:5.8S rDNA copy number ratio with increasing age at draw in the first blood draws of both the females and the males, and the bottom panel shows that this trend toward higher 18S:5.8S ratios at older ages is even more pronounced in the second blood draws, reaching statistical significance in the males (*P* = 0.00152).

**Fig 4.**
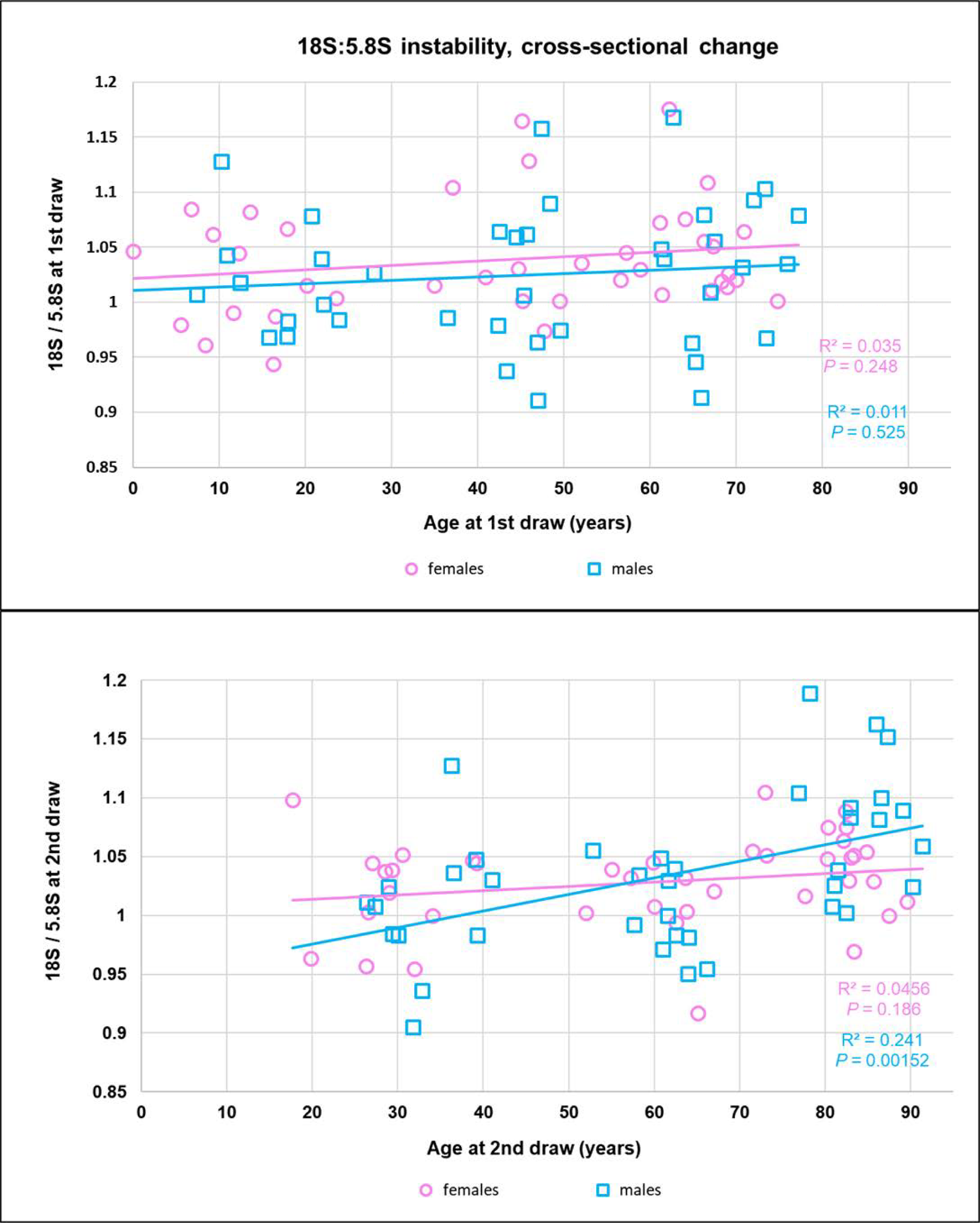
Cross-sectional 18S/5.8S rDNA ratios in the first blood draw DNAs, top panel, and in the second blood draw DNAs, bottom panel. Females’ datapoints are pink circles. Males’ datapoints are blue squares.

### Instability of the 18S:5.8S rDNA copy number ratio in longitudinal data

Fig 5 plots age at first blood draw on the x axis vs. (18S:5.8S at second draw) / (18S:5.8S at first draw) on the y axis. Both within-individual longitudinal increases and decreases in the 18S:5.8S ratio were observed across the lifespan, with increases at second draw occurring more often than decreases, in both sexes, and increases more likely at the oldest ages, particularly in the males. While there was no overall trend across the lifespan towards either an increasing or decreasing longitudinal change in the 18S:5.8S ratio in the females, an increasing longitudinal change in this ratio with age was seen in the males, which reached statistical significance (*P* = 0.0243).

**Fig 5.**
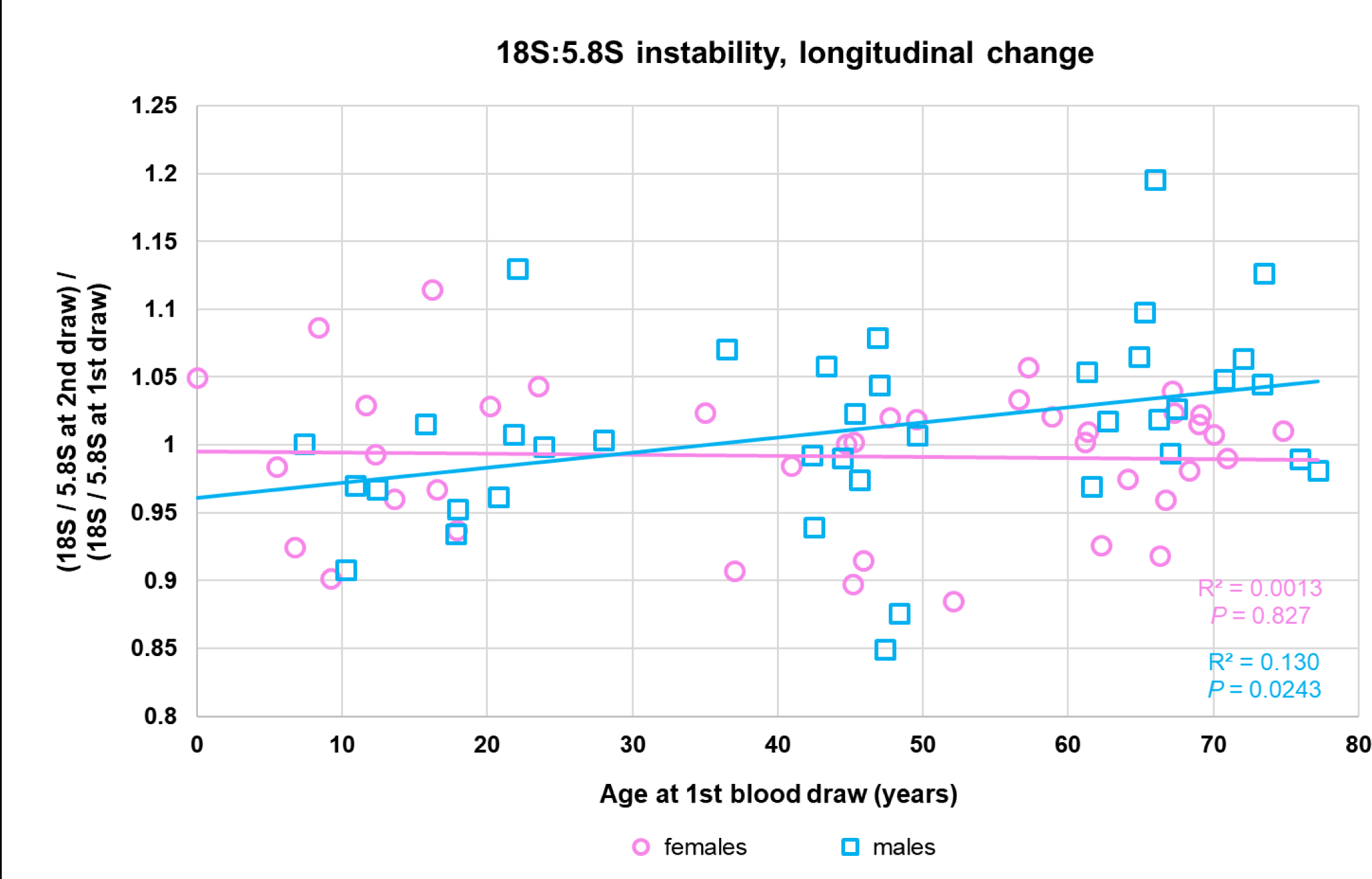
Instability of the 18S:5.8S rDNA ratio in longitudinally-collected blood samples. The 18S:5.8S ratio at second draw, relative to that at first draw, increased significantly across the lifespan in males, but not in females. Females’ datapoints are pink circles. Males’ datapoints are blue squares.

## Discussion

To our knowledge this study is the first to demonstrate rDNA copy number changes longitudinally across the lifespan in humans. Measures of rDNA instability may reflect the level of genomic instability generally. Cells with high rDNA instability (or high genomic instability generally) may be prone to undergoing cellular senescence or apoptosis, processes believed to contribute to the dysfunction, atrophy, and degeneration of various tissues with age. Alternatively, depending on the cell type and other conditions present, the rDNA (or general genomic) instability may contribute to the development of various cancers. These two novel MMqPCR assays of the 18S and 5.8S rDNAs will facilitate further studies to explore these possibilities. Recent studies in mice and humans have shown that the copy numbers of the 5S rDNA tandem repeats on Chromosome 1 approximately co-vary with the copy numbers of the 45S rDNA tandem repeats on Chromosomes 13, 14, 15, 21, and 22 [9]. Ribosomal DNA copy numbers in humans also appear to be coordinately regulated with the copy numbers of the mitochondrial DNA (mtDNA) [8]. Therefore, there may be healthy, as well as pathologic, ranges for the absolute and relative copy numbers of 5S rDNA, 18S and 5.8S rDNAs, and mtDNA. Further research will be needed to determine whether these quantitative traits will be clinically useful biomarkers of aging, accounting for a significant fraction of the variation in age at death in late middle-aged and older adults, and whether medical interventions stabilizing them to healthy ranges will be effective in extending the human healthspan.

### Human subjects, materials, and methods

#### Human DNA samples

The 47 three-generation Utah CEPH (Centre d’Etude du Polymorphisme Humain) families were originally contacted in the early 1980s for the purpose of building the first comprehensive human genetic linkage map. In the late 1990s to early 2000s many of the original Utah CEPH cohort were re-contacted for the Utah Genetic Reference Project (UGRP), in which new blood samples were drawn, and a large number of physical and biochemical traits were measured, with the goal of discovering heritable genetic variants influencing those traits [10]. The present study examines pairs of DNA samples longitudinally-collected from 40 females and 39 males from these families (25 from the youngest generation, 22 from the middle generation, and 32 from the grandparent generation), selected so that within each generation, all individuals of the same sex were unrelated. The first blood draw DNAs were extracted by standard phenol-chloroform methods. The second blood draw DNAs were extracted using the Gentra Puregene Blood Kit [11]. All extracted DNAs were dissolved in 10 mM Tris-Cl, 0.1-1 mM EDTA, pH 7.5 at 25°C, at a concentration of 100-200 ng/μl, confirmed by agarose gel electrophoresis to consist of high molecular weight DNA with negligible degradation, and stored long term at 4°C. The University of Utah’s Institutional Review Board has approved this research project.

#### Quantitative PCR

We used the monochrome multiplex polymerase chain reaction (MMqPCR) [12] with SYBR Green I as the detecting dye, to measure the rDNA copy number to single copy gene copy number ratio, relative to that in a reference DNA sample, by the Standard Curve method. The reference DNA was a pooled sample containing equal amounts of DNA from eight healthy individuals aged 65 years or older from the Utah population. The qPCR reaction mix composition used here is the same as in ref. 12, except for the primers used and the inclusion here of a restriction enzyme.

All primer sequences are written 5’ ‐> 3’:

MMqPCR 1: 18S rDNA primers: ATGGTAGTCGCCGTGCCT (final concentration, 1μM) and CGTCACCCGTGGTCACCAT (f.c. 1.5μM); beta-globin gene primers (with GC-clamps): CGGCGGCGGGCGGCGCGGGCTGGGCGGCTTCATCCACGTTCACCTTG (f.c. 500nM) and GCCCGGCCCGCCGCGCCCGTCCCGCCGGAGGAGAAGTCTGCCGTT (f.c. 500nM)

MMqPCR 2: 5.8S rDNA primers: CCTCGTACGACTCTTAGCGGT (f.c. 1μM) and GCACGAGCCGAGTGATCC (f.c. 1.5μM); albumin gene primers (with GC-clamps):

CGGCGGCGGGCGGCGCGGGCTGGGCGGAAATGCTGCACAGAATCCTTG (f.c. 400nM) and GCCCGGCCCGCCGCGCCCGTCCCGCCG GCATGGTCGCCTGTTCAC (f.c. 400nM)

The 18S rDNA to beta-globin gene relative ratios were measured in triplicate for each sample in one set of reactions, and the 5.8S rDNA to albumin gene relative ratios were measured in triplicate for each sample in another set of reactions. The Standard Curve reactions contained a 3-fold serial dilution of the reference DNA, with final input DNA amounts per reaction of 40 ng, 13.3 ng, 4.44 ng, 1.48 ng, and 0.494 ng of DNA.

TaqI restriction endonuclease (New England Biolabs) was included in the master mix to a final concentration of 4 units of TaqI per 10 microliter final PCR reaction volume. We found that restriction enzyme digestion of the genomic DNA prior to PCR is essential in order to obtain reproducible, reliable results in rDNA quantification.

The thermal profile for the 18S rDNA + beta-globin reaction was: 65°C x 60 min (TaqI digestion); 95°C x 15 min (DNA polymerase activation and DNA denaturation); and 40 cycles of 96°C x 15 sec, 58°C x 30 sec, 72°C x 15 sec with signal acquisition, 86°C x 15 sec with signal acquisition. The thermal profile for the 5.8S rDNA + albumin reaction was the same as above, except that the second signal acquisition was at 85°C instead of 86°C. All qPCR reactions were performed on a Bio-Rad CFX384 Real-Time PCR Detection System. Relative quantification was performed by the Standard Curve method using Bio-Rad’s CFX Manager Software Version 3.1. All statistical analyses were performed using the CFX Manager Software and Microsoft Excel.

Fig 6 shows Standard Curves for 18S rDNA amplification (top panel) and HBB amplification (bottom panel) for MMqPCR 1. Fig 7 shows Standard Curves for 5.8S rDNA amplification (top panel) and HBB amplification (bottom panel) for MMqPCR 2.

**Fig 6.**
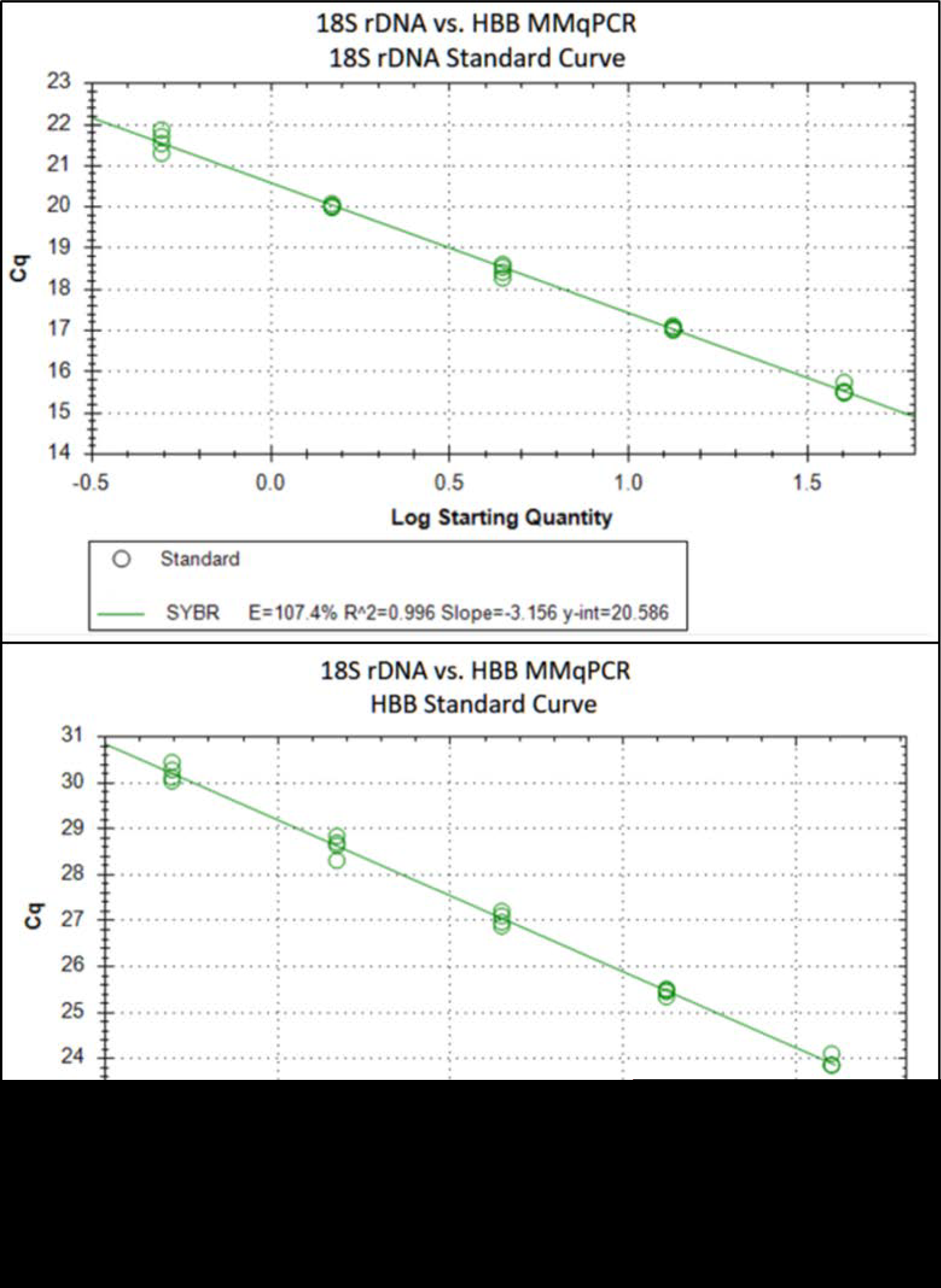
Standard curves for the 18S rDNA and the HBB (beta-globin gene) monochrome multiplex qPCRs. All subjects’ DNA samples were assayed in triplicate. The geometric mean of the coefficient of variation was 4.80% for the 18S/HBB ratios measured in the first blood draw DNAs, and 4.06% for the 18S/HBB ratios measured in the second blood draw DNAs.

**Fig 7.**
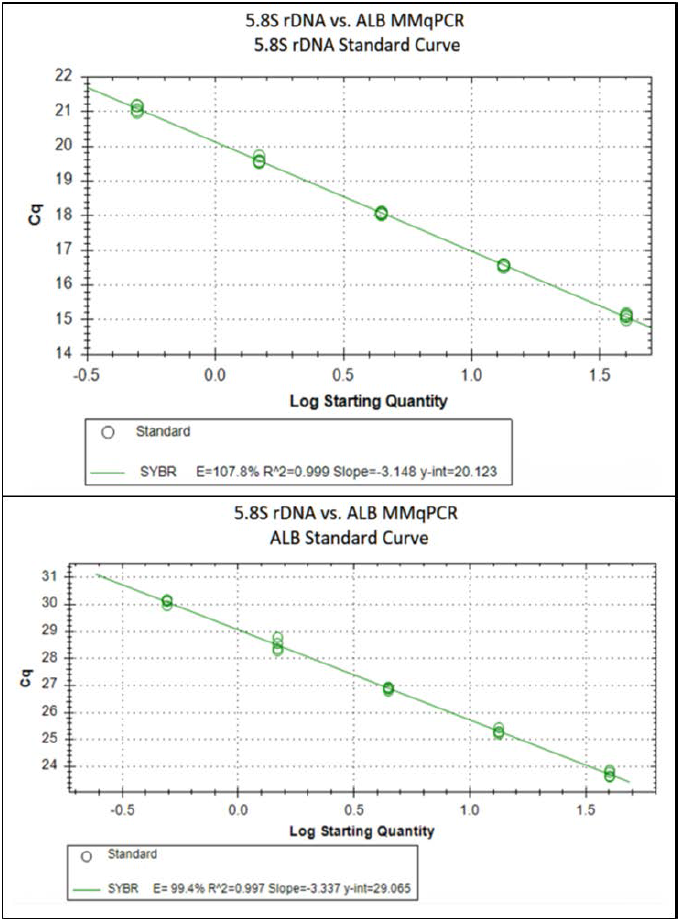
Standard curves for the 5.8S rDNA and the ALB (albumin gene) monochrome multiplex qPCRs. All subjects’ DNA samples were assayed in triplicate. The geometric mean of the coefficient of variation was 3.93% for the 5.8S/ALB ratios measured in the first blood draw DNAs, and 3.90% for the 5.8S/ALB ratios measured in the second blood draw DNAs.

## Data availability statement

Figures 1-7 have associated raw data collected during qPCR runs on Bio-Rad CFX384 Real-time Detection System, under the control of the Bio-Rad CFX Manager Software Version 3.1, and stored in computer files. Copies of these files are available, without restriction, from Dr. Cawthon, upon request at. All other relevant data are within the body of the paper and its figures.

## Acknowledgments

We thank all CEPH family members who participated in the Utah Genetic Reference Project; Andreas P. Peiffer, Medical Director of the UGRP; and Melissa M. Dixon, UGRP Study Coordinator.

## Author contributions

**Conceptualization**: Chris B. Ligerman, Richard M. Cawthon

**Data curation**: Chris B. Ligerman, Lisa Baird, Mark F. Leppert, Richard M. Cawthon

**Formal analysis**: Chris B. Ligerman, Richard M. Cawthon

**Funding acquisition**: Mark F. Leppert, Richard M. Cawthon

**Investigation**: Chris B. Ligerman, Richard M. Cawthon

**Methodology**: Richard M. Cawthon

**Project administration**: Chris B. Ligerman, Lisa Baird, Mark F. Leppert, Richard M. Cawthon

**Resources**: Lisa Baird, Mark F. Leppert, Richard M. Cawthon

**Supervision**: Richard M. Cawthon

**Validation**: Chris B. Ligerman, Richard M. Cawthon

**Visualization**: Chris B. Ligerman, Richard M. Cawthon

**Writing - original draft preparation**: Richard M. Cawthon

**Writing - review & editing**: Chris B. Ligerman, Lisa Baird, Mark F. Leppert, Richard M. Cawthon

